# The biofuel producing legume tree, *Pongamia pinnata*, offers strong potential for carbon sequestration

**DOI:** 10.1101/2025.06.04.657758

**Authors:** A Chen, P Scott, PM Gresshoff, BJ Ferguson

**Author notes:** Terragen Biotech Pty Ltd, Coolum Beach QLD 4573.

## Abstract

*Pongamia pinnata* (Linn.) Pierre, also called *Millittia pinnata* or Indian beach, is a fast-growing legume tree that has gained considerable attention for its potential as a feedstock for biofuel. In this study, we assessed the growth characteristics and carbon sequestering potential of Pongamia grown under sub-tropical environments. At two different sites located in Queensland, Australia, Pongamia was found to produce considerable biomass, even when planted in nutrient-poor environments having low soil organic carbon. The seedlings planted were all derived directly from genetically diverse, yet related seed. Variability observed in growth appeared to be associated with the mother tree from which the seeds were sourced, underscoring the importance of planting elite genetic material for commercial purposes. Pongamia is known to be drought tolerant, with findings here demonstrating that even young seedlings have this characteristic. Collectively, our findings indicate that Pongamia can grow quickly, reaching a biomass of 13-19 kg over 3-4 years in our studies, and sequester high quantities of carbon, at 2.9 to 4.0 t of carbon per ha (assuming a tree density of ∼450/ha), even when planted in suboptimal growing conditions on marginal, nutrient-poor lands.

## INTRODUCTION

*Pongamia pinnata* is a fast-growing tropical legume tree found in areas of India, southern Asia and Australia. It belongs to the Fabaceae family and can acquire essential nitrogen for growth and development through its ability to interact with beneficial nitrogen-fixing soil bacteria, broadly called rhizobia (Biswas and Gresshoff, 2014). This provides the tree with a competitive growth advantage over non-legume species and offers advantages to soil improvement, ecological restoration efforts and agroforestry systems. Able to grow in a variety of soil conditions, including saline, acidic and degraded soils, and to withstand wide temperature ranges, periodic flooding and drought conditions, Pongamia is widely recognised as a hardy species suitable for growth in a range of environments (Sangwan et al., 2010; Degani et al., 2022). These resilient attributes make it suitable for cultivation on marginal and degraded lands that are not suitable for many food crops. The adaptability of Pongamia to marginal land is particularly important as it provides economic benefits to rural communities and enables plantations to be grown in areas not suitable for most conventional crops, avoiding the unwanted displacement of food for fuel production (Leksono et al., 2021). Solutions to combat climate change and enhance energy security require sustainable alternatives, with Pongamia representing one such option to further explore.

In recent years, Pongamia has garnered considerable attention for its potential as a sustainable source of biofuel, including both renewable diesel (RD) and sustainable aviation fuel (SAF) (Klein-Marcuschamer et al., 2013). Its seeds contain 30-40% oil, which is rich in oleic acid, making it ideal for biodiesel production (Pavithra et al., 2012). Trees reach maturity in four to five years, with elite varieties producing over 10,000 oil-rich seeds per year. The oil can be extracted through mechanical pressing or solvent extraction processes. Pongamia biodiesel closely matches petroleum diesel in performance, with good cold flow, high cetane numbers, low sulphur content, and significantly reduced emissions reported (Karmee and Chadha, 2005; Dwivedi and Sharma, 2014; Arul Nicholas et al., 2022). Additionally, by-products like seed cake and biomass offer further economic and environmental benefits through uses such as animal feed, fertiliser, and biogas production (Degani et al., 2022).

Being a fast-growing tree, Pongamia also offers the additional benefit of storing large amounts of carbon. Carbon sequestration has been proposed as a critical process in mitigating climate change, involving the capture and storage of carbon dioxide (CO₂) from the atmosphere (Arul Nicholas et al., 2022). Through the process of photosynthesis, trees assimilate atmospheric CO₂ and store it in their biomass, including roots, stems, branches and leaves. The tree’s fast growth rate allows it to capture a significant amount of carbon in a relatively short period. Moreover, when leaves and other organic matter fall and decompose, their breakdown further contributes to the carbon-holding capacity of the soil.

Pongamia also has a robust root system that can reduce soil erosion, a major cause of soil carbon loss (Degani et al., 2022). By anchoring the soil and preventing erosion, Pongamia helps maintain the organic matter in the soil, which is essential for long-term carbon sequestration (Zeng et al., 2024). The trees also add considerable organic material to the soil through root exudates. This helps promote the population of soil microbes that largely rely on available carbon, further enhancing the soil organic carbon (SOC) content. Pongamia’s nitrogen-fixing capability (Samuel et al., 2013) also contributes to carbon sequestration by improving soil fertility and stability, making it more conducive to long-term carbon storage (Jensen et al., 2012). Moreover, improved nitrogen levels encourage greater plant growth in the surrounding area, leading to an increase in overall biomass and carbon uptake in the region (Chaukiyal et al., 2000; Jensen et al., 2012). Furthermore, nitrogen enrichment in the soil supports carbon stabilisation, as it promotes the formation of stable organic compounds that are less prone to decomposition.

This ability to remove carbon from the atmosphere and sequester it in both biomass and soil plays a pivotal role in offsetting greenhouse gas emissions (Jensen et al., 2012; Crutzen et al., 2016). Indeed, the significance of carbon sequestration in Pongamia extends beyond individual trees; it has implications for larger ecosystems and climate strategies. By integrating Pongamia into agroforestry programmes (Wani et al., 2009; Prasad, 2022), farmers can create multi-functional landscapes that provide economic, environmental, and social benefits (Sharma et al., 2016). These systems can enhance biodiversity, improve water quality, and support local economies through the production of biofuels and other tree-derived products (Sharma et al., 2016).

Preliminary research and growth observation indicate that Pongamia can sequester significant amounts of carbon over its lifespan. Estimates suggest that a mature Pongamia tree can store on average 20-30 kilograms of CO₂ per year depending on growth conditions, contributing to substantial reductions in atmospheric carbon levels when planted at scale (Wylie et al., 2023). This rate of carbon sequestration makes Pongamia a potentially viable option for carbon offset projects, especially in regions where marginal land is available. Effective management practices and appropriate planting strategies will be crucial to optimise this process and maximize these benefits (Degani et al., 2022). Understanding the specific growth conditions, optimal planting densities, and maintenance requirements will be essential for leveraging Pongamia’s full potential in carbon sequestration efforts.

As the world increasingly turns to carbon markets to reduce emissions, Pongamia plantations could be used to generate carbon credits that offset emissions (Degani et al., 2022). Indeed, Pongamia’s hardiness and adaptability, coupled with its strong potential for carbon capture, make it appealing for long-term carbon sequestration projects and potential for generating carbon credits (D’silva, 2003; Degani et al., 2022). Carbon markets have evolved to be one of the key mechanisms for reducing greenhouse gas emissions, representing a shift towards more sustainable practices, supporting climate action, and creating economic opportunities (Mitra et al., 2021). However, despite its considerable appeal, the carbon sequestration potential of Pongamia requires further research, largely due to the lack of existing biomass data available. This could be supported through a crediting method under carbon credit schemes for the Pongamia species using a ‘measured approach’ while data collection is ongoing, de-risking investment for those looking to establish biofuel feedstocks.

Here, we report on our investigations into the carbon sequestering potential of Pongamia. Plantations located in Queensland, Australia, provided opportunities to quantify and model the carbon sequestration potential of trees grown in different environments and with different management regimes. Findings indicate Pongamia is well adept to sequester large quantities of carbon, including when grown in substandard conditions on marginal land.

## RESULTS

### Spring Gully Pongamia trial

To investigate the growth and development of Pongamia trees differing in their genetics, and to understand their potential for carbon sequestration, a trial was planted in collaboration with BPA Pty Ltd in 2008 at Spring Gully, Queensland, Australia (land owned then by Origin Energy Australia; Figure 1). This location was selected for its marginal growing conditions, with a total of 680 trees planted in 20 rows on one and a half hectares of land (-26.003909, 149.077883). Spring Gully has a semi-arid climate, typically characterised by hot summers (30-35°C), mild winters (15-20°C) and low to moderate rainfall (Figure 1). Winter nighttime temperatures occasionally drop below 0°C but are typically around 5-10°C which can impact Pongamia growth (as air temperature below 17°C induce growth dormancy as seen by a lack of apical growth). Annual precipitation is approximately 500-700 mm, with the majority occurring during the summer months. The dry season occurs during winter, when rainfall is sparse, and soil moisture levels drop.

**Figure 1.**
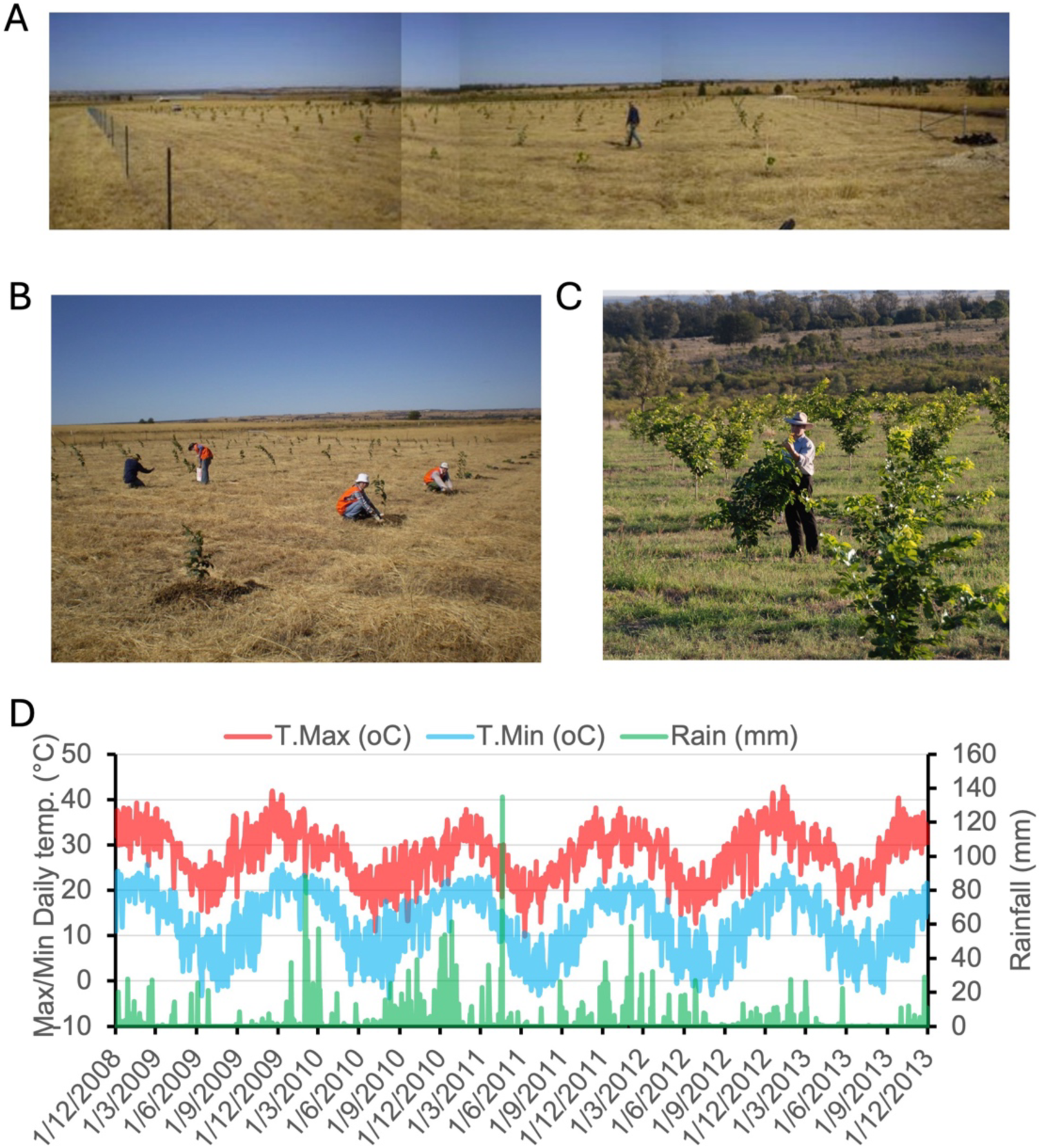
Pongamia plantation located at Spring Gully, Queensland, Australia. The plantation consists of 20 rows with 34 trees per row, located on 1.5 ha of land, managed as part of a 300 ha Pongamia plantation **(A, B)** Pongamia seedlings being planted at the site. **(C)** One year after planting, the seedlings are already showing robust growth. **(D)** Temperature and rain fluctuation patterns at Spring Gully (-26.003909, 149.077883), visualised in 3-month intervals. Minimum (blue) and maximum (red) daily temperature in °C are plotted. Average daily rainfall in mm were plotted on a separate axis. All weather data obtained from https://www.longpaddock.qld.gov.au/silo/point-data/.

Seedlings from six elite mother trees were germinated from seed and maintained in a glasshouse until reaching a suitable size for planting in the field (Supplemental Table 1). At planting, basal diameter of the trunk and plant height were measured, with variation associated with their genetics being evident (Figure 2A-B). One-year after planting, the average basal diameter of the primary tree trunk located 0 – 2.5 cm above ground was 2-3 cm (Figure 2C). By 3.5-years after planting, the diameter was 4-6 cm (Figure 2C). Mature Pongamia trees anticipated to be suitable for commercial scale harvest are often reported to have a trunk diameter of 20-50 cm, allowing for manageable maintenance and harvest of the trees (Bohre et al., 2014).

**Figure 2.**
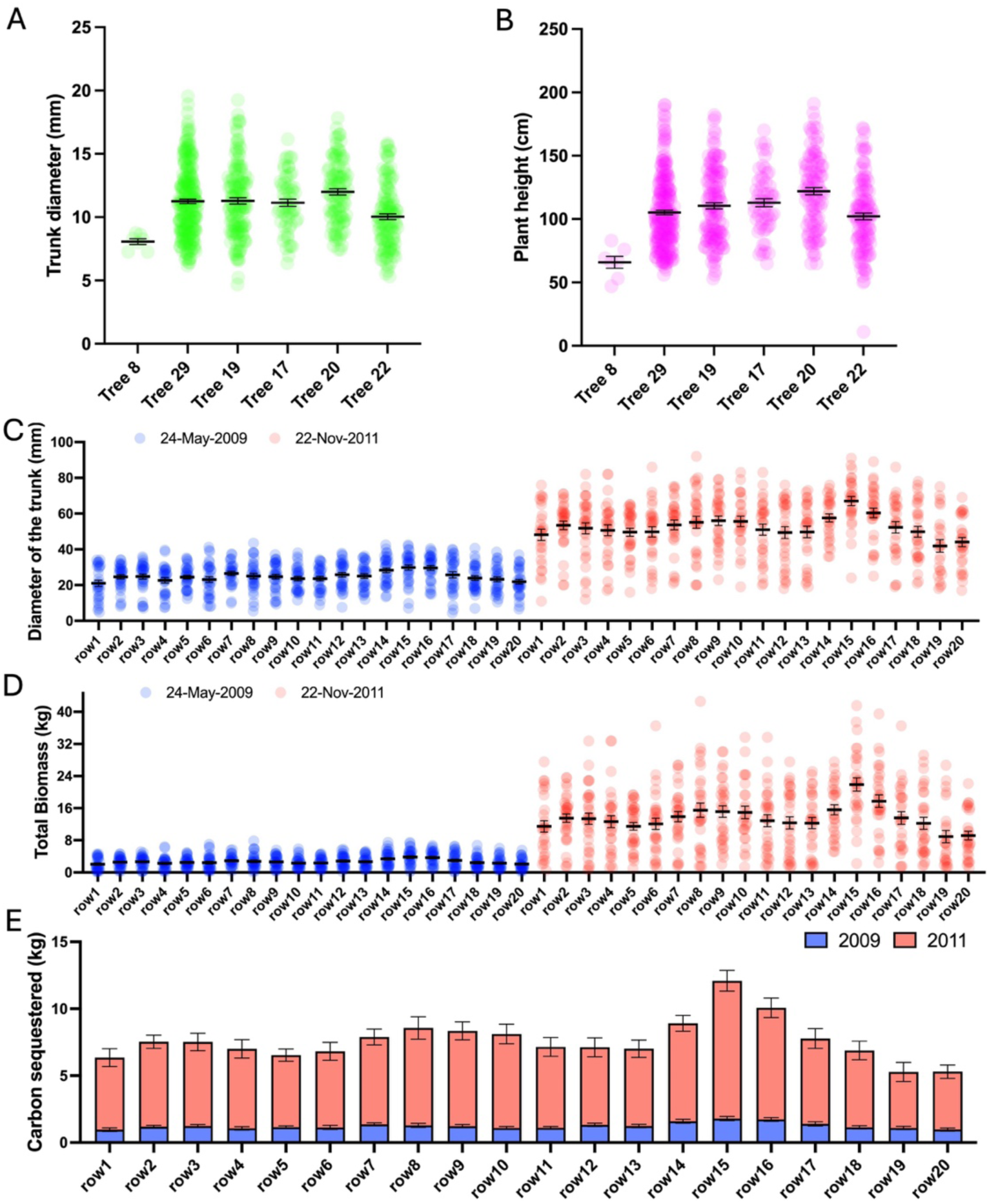
Growth and estimated carbon sequestered of Pongamia tress at the Spring Gully plantation. **(A)** Plant height and **(B)** weight of the various Pongamia genotypes at time of planting (listed in Supplemental Table 1). **(C)** Trunk diameter (*i.e.,* basal diameter) of trees measured at a 0 – 2.5 cm height above ground for a total of 680 trees evenly planted in 20 rows. The trees were measured at 1 (24-May-2009, blue dots) and 3.5 (22-Nov-2011, red dots) years after planting. **(D)** Above- and below-ground biomass of all trees at the plantation. **(E)** Average amount of carbon sequestered per tree per row. Error bars represent the standard errors of the mean.

Allometric equations are commonly used to estimate biomass based on measurable variables such as trunk diameter and plant height. Total biomass was predicted here using the allometric equation 0.609*EXP(-0.667+1.784*LN(d)+0.207*(LN(d))^2-0.0281*(LN(d))^3) (Chave et al., 2005). Specific gravity of 0.609 was derived from Rajput et al. (1985) and below-ground biomass predicted using 0.275*above-ground biomass (Figure 2D) (Mokany et al., 2006). Dry biomass estimates were converted to carbon using the default carbon fraction of 0.47 kg carbon per kilogram of dry weight (recommended by IPCC Guidelines for National Greenhouse Gas Inventories). Biomass carbon was converted to CO_2_ using 44/12, the molecular weight ratio of CO_2_/C. Carbon sequestration estimates per row (34 trees) indicates high variation between individual trees (Figure 2E). This is most likely attributed to genetic differences caused by the outbreeding nature of Pongamia, as environmental parameters did not vary widely. In subsequent years after planting, variation continued to be detected in tree height and trunk diameter of trees from different parents, further highlighting the impact of genetics on development (Figure 2A-B, Supplemental Table 1). Those from Pongamia stock ‘Tree 20’ were estimated to sequester the most carbon of the genetic stocks planted (Figure 2). Overall, by 3.5 years post-planting, the trees had an average total biomass of 13.56 kg per tree and the trial had sequestered an estimated 2.89 t C/ha (10.59 t CO_2_/ha) at a planting density of 453 trees per hectare.

Interestingly, in September 2009, following the winter dormancy period, a late frost occurred after the trees had developed new leaves. Frost damage was observed as blackening and abscission of leaves. Some trees did not survive, but the majority did and fully recovered. Approximately 20 % suffered stem tip die-back (10–30 cm), which did not affect growth in subsequent seasons. These observations indicate that Pongamia can withstand mild frost damage, consistent with previous reports (Prasad and Pandey, 1987). This further demonstrates the hardiness of the trees.

### Analysis of Pongamia yield at Spring Gully in 2021

Visual inspection of the trees in 2021, 13.5 years after planting, found their size and growth to be relatively uniform (Figure 3A-B). Emergence of new leaves were abundantly observed on trees, indicative of vigorous growth (Figure 3C). Seeds collected from freshly harvested pods are medium to large, about 1-2 cm in length and 0.5-1.5 cm in width (Figure 3D). Seeds are typically oval or elliptical, with a smooth, slightly shiny surface, and weigh about 1-4 grams. Total pod harvesting was performed on three individual trees; two receiving irrigation and the other being unirrigated and relying on natural rainfall. The irrigated trees produced over 33 kg of pods per tree with an average seed weight of 2.2 to 2.75 g, whereas the unirrigated tree produced 28 kg of pods with an average seed weight of only 1.5 g. Interestingly and unexpectedly, the irrigated trees had approximately 6,000 to 7,500 seeds harvested, whereas 9,333 seeds were harvested from the unirrigated tree. This may indicate that irrigation had a negative effect on seed set and positive effect on seed weight, but could also be a random effect based on low sample size. More research using different genetic varieties grown across different locations is needed to fully understand the impact of irrigation on yield.

**Figure 3.**
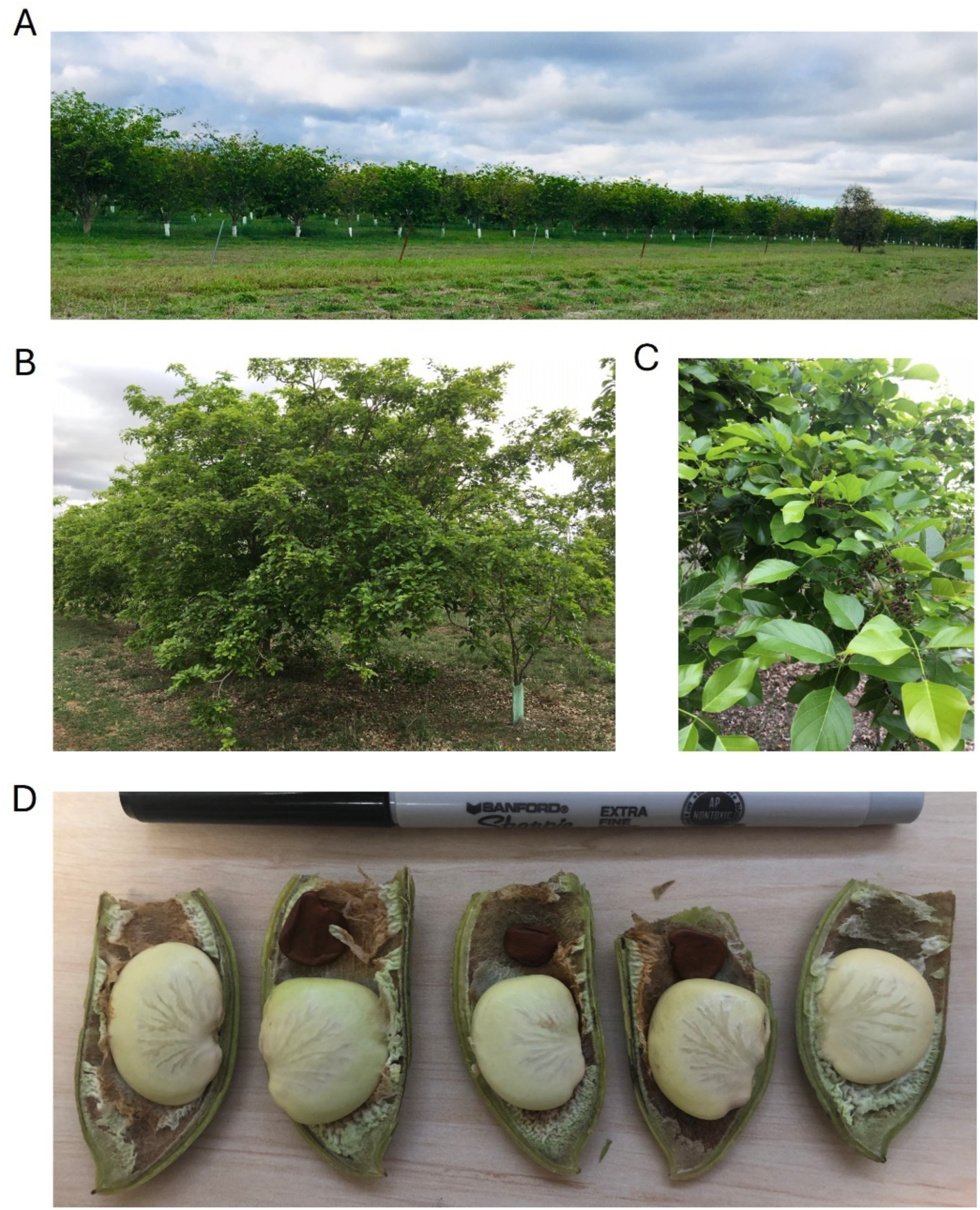
Spring Gully Pongamia plantation 13.5 years after planting. **(A)** A row of Pongamia plants irrigated with treated wastewater. **(B)** An individual tree exhibiting an extended globose canopy with an intensive branching pattern. **(C)** Emergence of new leaves and flower bunches indicative of vigorous growth. **(D)** Pods containing near-mature seeds collected from an irrigated tree. Note the seed uniformity and the aborted second seed (seen in the three central pods).

It is broadly known that tree biomass and seed yield are influenced by water availability, with water stress significantly affecting growth, flowering and pod formation. Despite Pongamia being highly drought-tolerant and able to survive in areas with annual rainfall as low as 500 mm, trees under drought-conditions typically exhibit reduced growth, impacting their carbon sequestration potential, and may produce fewer flowers and pods, and/or smaller seed, impacting oil yield (Degani et al., 2022). Optimal growth for Pongamia carbon storage and pod formation requires annual rainfall of around 800-1200 mm in most regions (Degani et al., 2022). Our findings highlight the need for more research in this area, including how supplemental irrigation could impact carbon sequestration and yield.

### Gatton Pongamia trial

A Pongamia trial was established in 2008 at the University or Queensland, Gatton research fields in Gatton, Queensland, Australia (-27.54020, 152.33471). This trial aimed to further assess the growth, development and carbon sequestration potential of trees. Unlike Spring Gully, the Gatton location represents a high-quality growing region for the trees. Young seedlings derived mainly from seed of various genetic backgrounds were established in a glasshouse and transferred to the field in rows (Figure 4). Temperature profiles typically had a maximum of 20-35°C between 2008 to 2013 (Figure 4). Minimum temperatures during winters reached as low as 0-5°C from June to September. Winters were relatively dry, with the majority of rainfall occurring in summer, reaching as high as 110 mm in the first quarter of 2013 (Figure 4).

**Figure 4.**
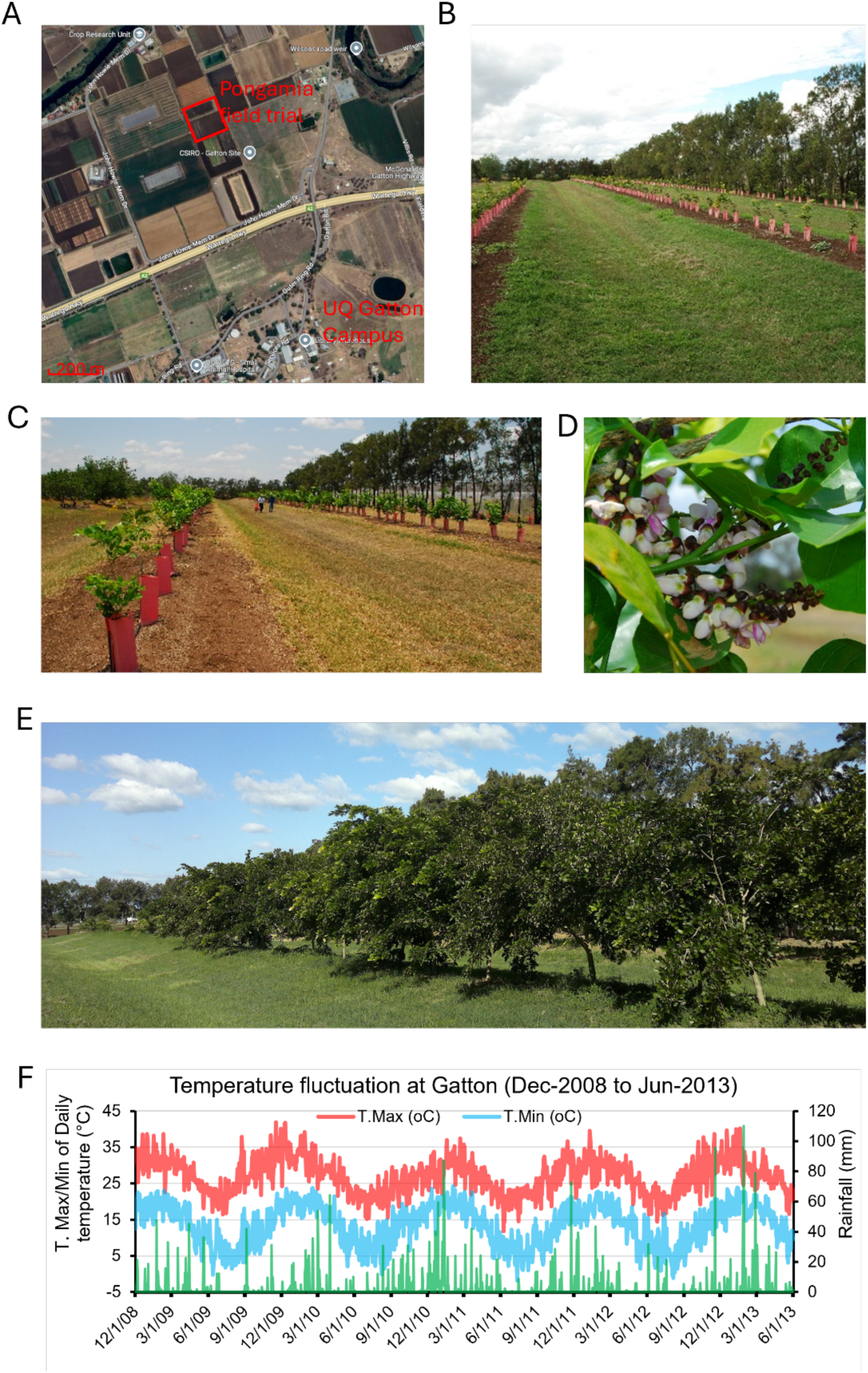
Pongamia field trial at Gatton, Queensland. **(A)** Satellite image of the Pongamia field trial at the University of Queensland, Gatton campus (27.54020, 152.33471). Highlighted in a red box is the location of the trial area. **(B)** Seedlings following planting in 2008. **(C)** Trees growing one year after planting in January 2009. **(D)** Flowers forming on some of the trees one year after planting. **(E)** Trees growing five years after planting in March 2013. **(F)** Temperature fluctuation at Gatton visualised in 3-month intervals. Minimum (blue) and maximum (red) daily temperature in °C are plotted. Average daily rainfall in mm are plotted on a separate axis. All weather data obtained from https://www.longpaddock.qld.gov.au/silo/point-data/.

As they matured, the trees at Gatton exhibited uniform height and a globose canopy structure (Figure 5). Total biomass consisting of both above- and below-ground parts, was quantified at 12 time points during the period between December 2008 to June 2013 (Figure 5). Significant biomass increases were not evident within the first months of growth, but quickly accelerated thereafter (Figure 5). Strong variability in tree size also became apparent with age (Figure 5). This is largely considered to be due to differences in genetics between trees and underlines the importance of planting good genetic stock in plantations. As a result, some trees grow faster and establish better than others. Within 15 months, the trees grew as tall as 3 m in height and up to 2 kg in above-ground dried biomass, with 6% exhibiting seed-bearing pods. Within 54 months, roots were established in deep soil layers (estimated at about 2-3 meters depth), enabling trees to tolerate periods of water deficit without signs of wilting or stress, likely due to the roots accessing the water table. Assuming the carbon content of the above-ground biomass was 47% of the total dry weight (Chave et al., 2005), an average of 19.3 kg total biomass per tree at 4.5 years equates to 9.07 kg carbon per tree. At a plantation density of 450 trees per hectare, this equates to approximately 4.08 t C/ha (14.96 t CO_2_/ha) sequestered.

**Figure 5.**
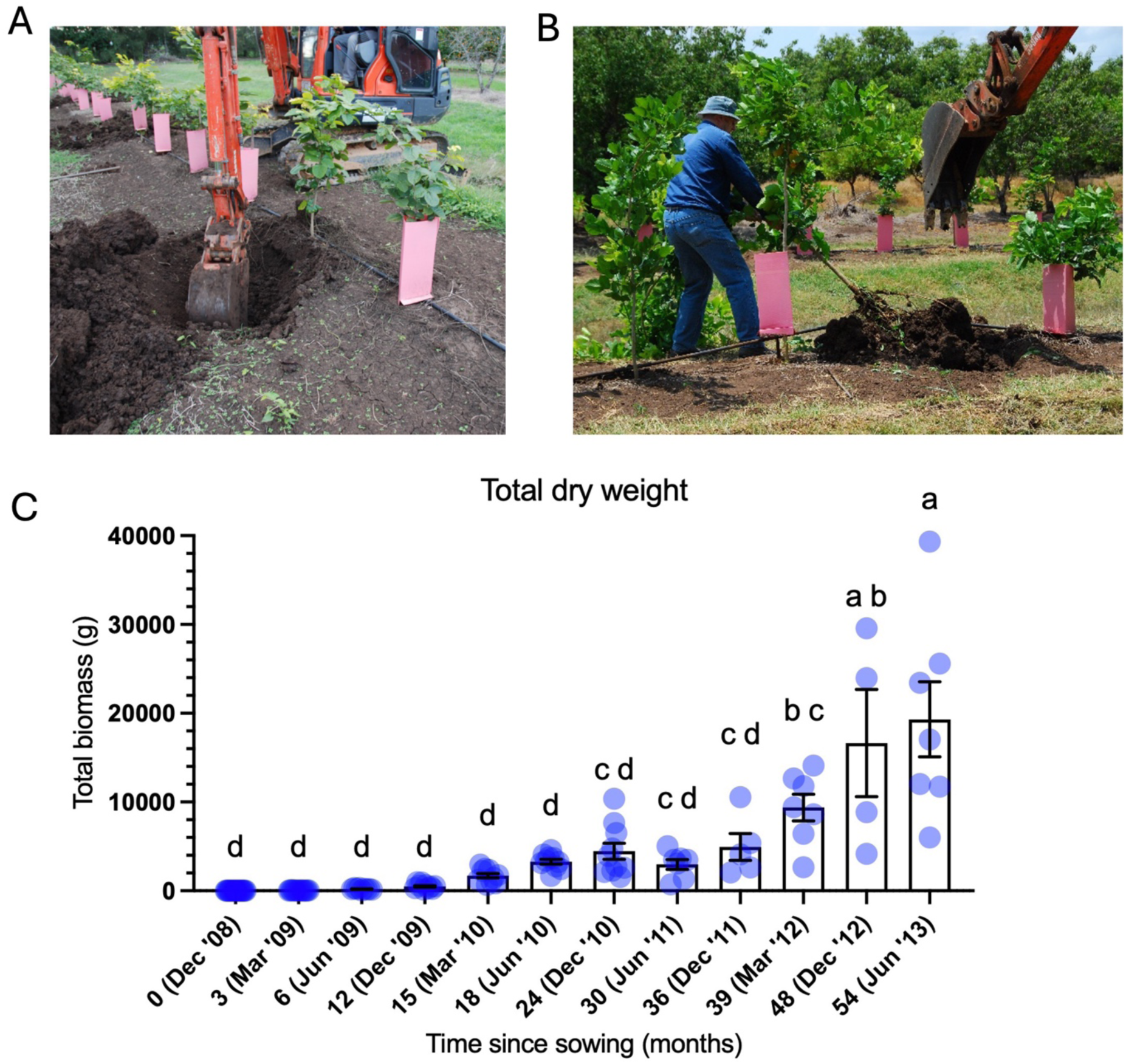
Biomass of Pongamia trees from the Gatton field trial. **(A)** An excavator was used to help uproot trees and access their root system biomass. **(B)** Uprooting a tree to assess root and shoot biomass one year after planting in January 2009. **(C)** Total above- and below-ground biomass accumulation over 54 months. Above-ground biomass was determined by directly weighing the material following drying. Below-ground biomass within the first year after planting was also measured directly, after which they were estimated using the formula 0.275*biomass (above-ground). Means and standard error of the mean are plotted. One-way ANOVA was performed with a post-hoc Tukey test, with different letters indicating significant differences between samples (p ≤ 0.05). A harmonic mean sample size of 7.99 was used to account for unequal variance across samples.

### Growth Characteristics of Trees from the Gatton Pongamia trial

Early assessment of Pongamia tree growth at the Gatton plantation 0-12 months post-planting enabled growth characteristics to be modelled as a sigmoidal curve using the Pade approximant on shoot dry weight and shoot length (Figure 6). An initial slow growth phase was identified which likely resembled a lag as the roots established themselves and the seedlings recovered from the shock of being planted. This was followed by an intermediate phase where rapid growth was observed over the summer followed by a plateauing phase associated with the cooler months. This growth pattern is similar to other reports assessing Pongamia growth following planting (Goudriaan and Van Laar, 2012; Dambreville et al., 2015).

**Figure 6.**
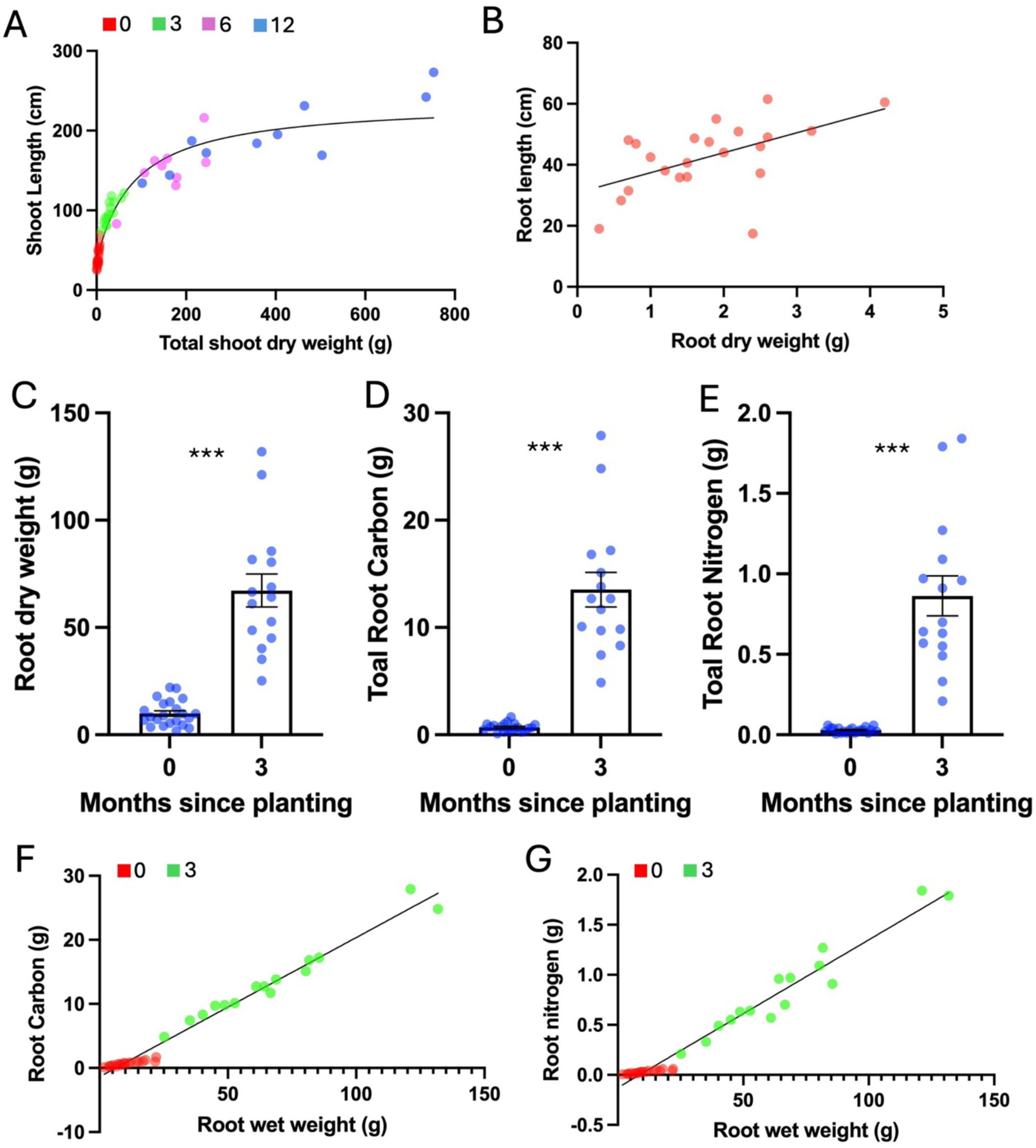
Growth characteristics of Pongamia trees grown at Gatton, QLD. **(A)** Shoot dry weight was correlated with shoot length. Regression was modelled using Pade (1,1) approximant, Y=(A0 + A1*X)/(1 + B1*X). R squared was 0.9269. Data were collected at 0 (red), 3 (green), 6 (magenta) and 12 (blue) months post planting. **(B)** Root length was correlated with root dry weight. Linear regression was modelled on Root length in centimetres to root dry weight in grams at planting (0 months). R squared was 0.2917. Root length was difficult to measure reliably due to the challenge of extracting intact roots from soil. **(C-G)** Nitrogen and carbon accumulation in the roots of Pongamia during the first 3 months of growth at Gatton, QLD. **(C)** Total root dry weight in grams, **(D)** Carbon content in grams and **(E)** Nitrogen content in grams were determined at 0 and 3 months from the initial planting. Pair-wise t-test at p ≤ 0.05 was independently performed on 0 compared with 3 months for each of the total root dry weight, carbon, and nitrogen datasets. Means and standard error of the mean were plotted. *** indicates statistical significance at p ≤ 0.001 by one-way ANOVA. **(F)** Linear regression was modelled on root Carbon and **(G)** root Nitrogen against root fresh weight at 0 (red) and 3 (green) months post planting. A clear linear relationship was observed between root carbon content and root wet weight, with an R squared value of 0.9791. The equation was Y = 0.2171*X - 1.359. A clear linear relationship was also observed between the root nitrogen content and root wet weight, with an R squared value of 0.9653. The equation was Y = 0.01476*X - 0.1271.

Root dry weight was positively correlated with root length 3 months post-planting (Figure 6B). Total root biomass, nitrogen and carbon were also measured at 0- and 3-months post-planting. The increase in growth positively correlated with significant increases in carbon and nitrogen content of the roots at p = 0.05 (Figure 6C-E). On average, carbon content in Pongamia roots was about 40% of the total dry biomass (Figure 6D). The nitrogen content was about 2% of root dry weight (Figure 6E). Root carbon and nitrogen content showed a direct proportional relationship with the root wet weight of the trees. (Figure 6F-G). Some of the Pongamia seedlings start to flower and produce pods 12 months after planting. The total pod harvest per tree significantly increased over the subsequent four years with a harvest weight of 1 kg at 48 months, with the seeds representing approximately 50% of the total harvest.

### Soil analyses reveal differences between the Spring Gully and Gatton field sites

Soil organic carbon (SOC) levels markedly impact soil health and crop productivity. At the Gatton site, the SOC level was 1.26%, while at Spring Gully it is in the range of 0.36–1.5%, indicating greater limitations for cropping and biomass accumulation (Table 1). SOC content of 2% is deemed to be the threshold for productive cropping in temperate regions (Loveland and Webb, 2003), with levels below 1.1% reported to cause reductions in crop yield up to 20% (Aune and Lal, 1997). Hence, our analyses confirmed the soil at both sites was sub-optimal, particularly at Spring Gully, and the land would be considered marginal for production with standard crops. To address these challenges, site managers at Gatton employ practices such as the use of organic supplements and cover cropping to enhance SOC levels. These methods improve soil structure, water retention, and nutrient availability, supporting better plant growth. A targeted soil management strategy was not employed at Spring Gully, but would be of benefit to help optimise SOC and promote better productivity and ecological resilience.

**Table 1.** Nutrients and trace elements detected in soil samples collected from the Spring Gully and Gatton trial sites. TN = Total Nitrogen; SOC = Soil Organic Carbon, CEC = Cation Exchange Capacity; Extr S = Extractable Sulphur; cmol(+)/kg = centimoles of positive charge per kilogram of soil. PBI = Phosphorus Buffering Index; EC = Electrical Conductivity; meq = milliequivalents per 100 grams; CEC = Cation Exchange Capacity; DTPA = diethylenetriaminepentaacetic acid; Colwell P = Colwell P soil phosphorus test; Colwell K = Colwell K soil potassium test.

| Spring Gully | Conc. | Lower | Upper | Gatton | Conc. | Amount |
| --- | --- | --- | --- | --- | --- | --- |
| TN | Wt % | 0.049 | 0.126 | TN | Percentage | 0.12% |
| SOC | Wt % | 0.358 | 1.534 | SOC | Percentage | 1.26% |
| Ca | mg/kg | 4849 | 5095 | Ca (NH<sub>4</sub>Cl) | meq/100g | 10.68 |
| K | mg/kg | 183 | 244 | K (NH<sub>4</sub>Cl) | meq/100g | 0.8 |
| Mg | mg/kg | 684 | 850 | Mg (NH<sub>4</sub>Cl) | meq/100g | 9.77 |
| Na | mg/kg | 388 | 702 | Na (NH<sub>4</sub>Cl) | meq/100g | 0.47 |
| CEC | cmol(+)/kg | 32.13 | 35.93 | CEC | meq/100g | 21.72 |
| Ca | cmol(+)/kg | 24.19 | 25.41 | Ca | % CEC | 49.20% |
| K | cmol(+)/kg | 0.47 | 0.63 | K | % CEC | 3.68% |
| Mg | cmol(+)/kg | 5.63 | 7 | Mg | % CEC | 44.98% |
| Na | cmol(+)/kg | 1.69 | 3.05 | Na | % CEC | 2.16% |
| Colwell P | mg/kg | 4.8 | 12.8 | Colwell P | mg/kg | 64.2 |
| Extr S | mg/kg | 2.9 | 5.6 | S (KCl) | mg/kg | 17 |
| Cu | mg/kg | 1 | 2.3 | Cu (DTPA) | mg/kg | 0.82 |
| Fe | mg/kg | 12 | 17.1 | Fe (DTPA) | mg/kg | 45.88 |
| Mn | mg/kg | 1.8 | 9.6 | Mn (DTPA) | mg/kg | 15.5 |
| Zn | mg/kg | 0.5 | 1.6 | Zn (DTPA) | mg/kg | 1.57 |
| As | mg/kg | 0.03 | 0.13 | CaCl<sub>2</sub> | pH | 5.95 |
| Cd | mg/kg | 0.01 | 0.06 | Colwell K | mg/kg | 438.6 |
| Co | mg/kg | 0.01 | 0.06 | Cl | mg/kg | 52 |
| Cr | mg/kg | 0.02 | 0.03 | Nitrate-N (H<sub>2</sub>O) | mg/kg | 21.1 |
| Hg | mg/kg | 0.02 | 0.02 | EC | dS/m | 0.12 |
| Ni | mg/kg | 0.1 | 0.47 | Al (KCl) | meq/100g | 0.02 |
| Pb | mg/kg | 0.9 | 1.6 | PBI | units | 52.2 |
| Se | mg/kg | 0.1 | 0.2 | Boron | mg/kg | 0.65 |
|  |  |  |  | Ca:NO<sub>3</sub> ratio | ratio | 2.53 |
|  |  |  |  | Ca:Mg ratio | ratio | 1.09 |
|  |  |  |  | NO<sub>3</sub>-N (KCl) | mg/kg | 19.8 |
|  |  |  |  | NH<sub>4</sub>-N (KCl) | mg/kg | 7.9 |
|  |  |  |  | Total P | mg/kg | 1240 |
|  |  |  |  | H<sub>2</sub>O | pH | 6.72 |

Total nitrogen (TN) levels at Gatton and Spring Gully also reveal critical differences in their capacity to support productive cropping. At Gatton, TN was 0.12%, whereas at Spring Gully it ranges from 0.05% to 0.12% (Table 1). This is indicative of insufficient N for optimal tree growth. Being a legume capable of forming nitrogen-fixing root nodules, Pongamia can overcome issues of low soil N to a great extent, with additional strategies such as incorporating nitrogen-fixing cover crops or applying organic or synthetic nitrogen-based fertilisers also able to address N deficiencies to enhance crop performance.

At the Gatton site, the potassium level was 438.6 mg/kg (Colwell K test), which is adequate to support productive cropping. In contrast, potassium levels at Spring Gully ranged from 183 to 244 mg/kg (Table 1). This deficiency would adversely affect plant growth and productivity as potassium plays a vital role in plant-water regulation, photosynthesis, stress tolerance, and overall plant health, with its availability closely linked to maximising crop yields. To address this limitation at Spring Gully, soil amendments could be applied, with the incorporation of organic matter also helping to improve potassium retention and availability over time. Levels of additional nutrients, key elements and components of soil and water samples from the Gatton and Spring Gully field sites are also provided in Table 1 and Supplemental Table 2, further illustrating the suboptimal conditions at Spring Gully.

### Allometric Modelling of Pongamia Growth

The diameter at breast height (dbh) of Pongamia grown in the northern tropical forests of Singrauli, India, were reported for trees up to 18 years old (Bohre et al., 2014). A positive correlation between biomass and plant age was observed from randomly selected trees having a dbh of ≥ 5 cm (Bohre et al., 2014). Using this data, we were able to apply an allometric model for estimating biomass in multiple environments (dry, moist, wet, mangrove, all types; Chave et al., 2005) to assess the growth rate and biomass accumulation of these trees (Figure 7A). Rapid growth of the trees was observed after 10 years, with accelerated growth occurring from year 15-18 (Figure 7A). This demonstrates the fast growth and considerable carbon sequestering potential of Pongamia, and the need for strong management practices and maintenance regimes to maximise tree growth and optimise seed production.

**Figure 7.**
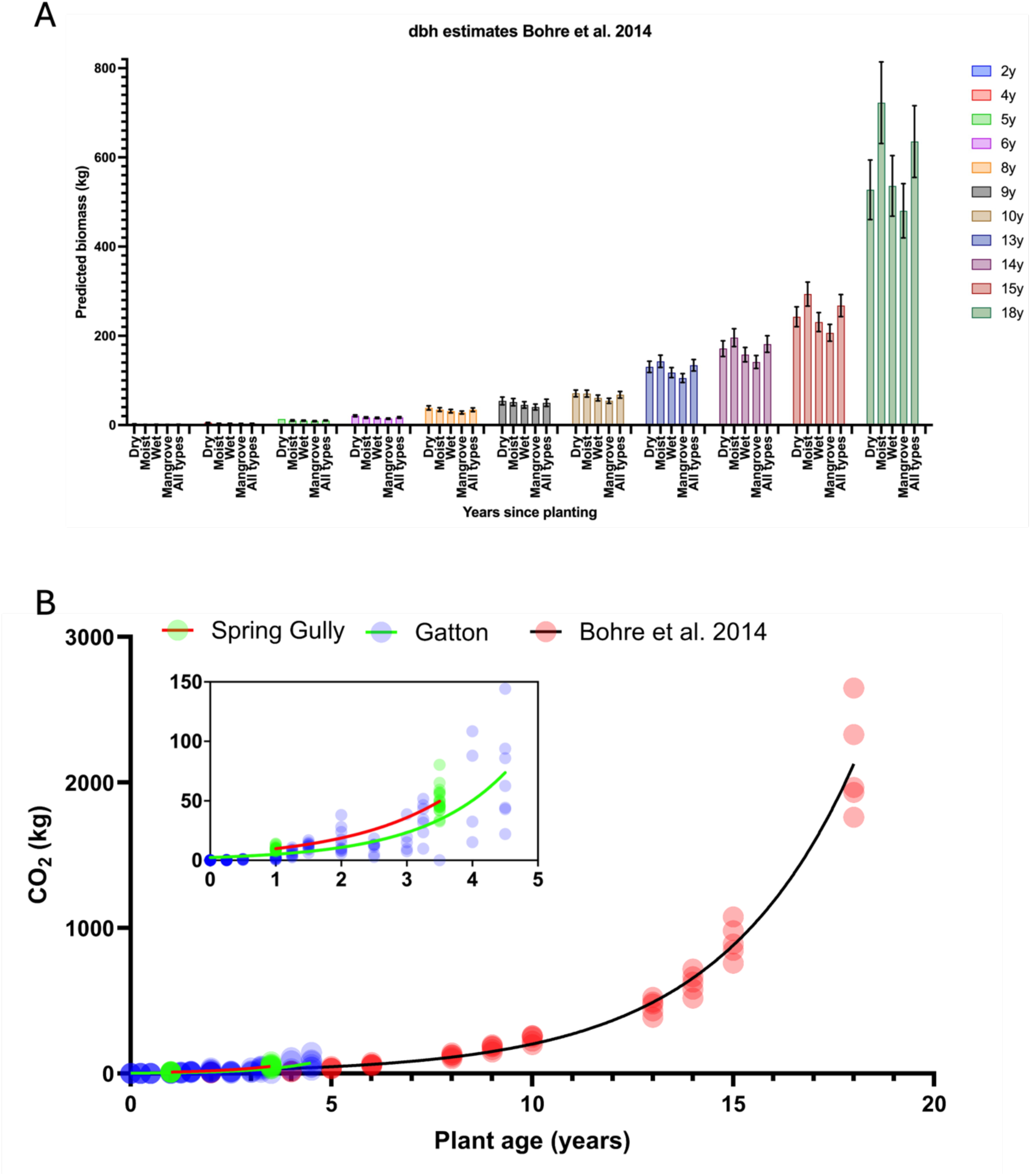
Estimated total biomass and CO_2_ sequestered of Pongamia trees from plantations in India and this study. **(A)** The diameter at breast height (dbh) of 2 - 18 year-old Pongamia trees grown in India (Bohre et al., 2014) was used to establish biomass. An allometric formula was used to estimate the above-ground biomass accumulated under dry (tropical), moist, wet, mangrove and all types of forest conditions (Chave et al., 2005). The below-ground biomass was estimated using the formula 0.275*biomass (above-ground). Error bars indicate standard deviations of the mean based on the dbh measurements. **(B)** Amount of CO_2_ sequestered per tree based on biomass accumulation illustrated using an exponential (Malthusian) growth model. Exponential growth was modelled as Y=Y0*exp(k*x), where Y is biomass, Y0 is the starting biomass, K is the Rate constant for growth, and x is time. Goodness of Fit showed an R squared value of 0.67 for the data from Gatton and 0.97 for the data derived from Bohre et al. (2014). Inset shows the amount of CO_2_ sequestered at Gatton and Spring Gully sites in Australia during the first five years of growth.

Biomass accumulation and CO_2_ sequestration of Pongamia trees planted at Gatton over 4.5 years can be modelled as exponential growth (Figure 7B, Supplemental Figure 1). The amount of CO_2_ sequestered per tree at Gatton follows a similar trajectory to the trees observed by Bohre et al. (2014) in Singrauli, and appears to be surpassing them at 4.5 years (Figure 7B, Supplemental Figure 1). Differences would be the result of classical factors relating to genetics, environment and management. Additional data collected from a range of plantations would provide more insight into this research area.

Collar diameter, crown width and tree height are considered reliable predictor variables for estimating total biomass of Pongamia trees using allometric equations (Rao et al., 2017), with predictive models for dry forests being increasingly accurate when using data from a large number of trees (Brown, 1997; Chave et al., 2005). A conservative estimate of carbon sequestered for the 4-year-old Pongamia plantation in Singrauli, India (Bohre et al., 2014) was 2.18 t C/ha (7.99 t CO_2_/ha). In this study, the cumulative data suggests that the total amount of carbon sequestered for the Spring Gully (3.5 years) and Gatton (4.5 years) trials was 2.89 and 4.08 t C/ha respectively, which equates to 10.59 and 14.96 t CO_2_/ha, respectively.

### Carbon sequestration potential of Pongamia pinnata

Estimated carbon sequestered by Pongamia and associated trees growing in tropical – subtropical regions is captured in Table 2. As anticipated, differences attributed to age are clearly evident as older individuals have sequestered considerably more carbon than young, establishing trees due to their size. Genetics, environment and management would also have an impact on tree growth. For example, Pongamia trees growing at Gatton, where conditions for growth are favourable, sequestered much more carbon (9.07 kg in 4.5 years) than trees growing at Spring Gully (6.37 kg in 3.5 years), where rainfall, temperature and soil conditions are all suboptimal for growth (Figures 1 and 4; Tables 1 and 2; Supplementary Tables 1). In Singrauli, 4-, 10-, and 18-year-old Pongamia trees (Bohre et al., 2014) sequestered an average of 2.4, 33.4, and 247.8 kg of carbon per tree, respectively. This equates to 8.8, 122.5, and 908.6 kg of CO_2_.

**Table 2.** Carbon sequestration potential of *Pongamia pinnata* and other tree species.

| Common name | Scientific name | Location | Number of trees assessed | Average trunk diameter* | Age of the trees | Above-ground biomass (kg)<sup>#</sup> | Total biomass (kg) | Carbon sequestered per tree (kg) | Reference |
| --- | --- | --- | --- | --- | --- | --- | --- | --- | --- |
| Pongamia | *Pongamia pinnata* | Spring Gully, QLD | 593<sup>^</sup> | 5.2 | 3.5 years | 10.63<sup>#</sup> | 13.56 | **6.37** | This study |
| Pongamia | *Pongamia pinnata* | Gatton, QLD | 7 | N/A | 4.5 years | 15.15<sup>#</sup> | 19.3 | **9.07** | This study |
| Ponagmia | *Pongamia pinnata* | St Lucia, QLD | 1 | 39.7 | 11 years | 902.6<sup>#</sup> | 1150.8 | **540.9** | This study |
| Pongamia | *Pongamia pinnata* | Singrauli, India | Unknown | 3.6 | 4 years | 4.1<sup>±</sup> | 5.2 | **2.4** | Bohre et al. 2014 |
| Pongamia | *Pongamia pinnata* | Singrauli, India | Unknown | 11.5 | 10 years | 55.7<sup>±</sup> | 71.0 | **33.4** | Bohre et al. 2014 |
| Pongamia | *Pongamia pinnata* | Singrauli, India | Unknown | 27.8 | 18 years | 413.4<sup>±</sup> | 527.1 | **247.8** | Bohre et al. 2014 |
| Madhushevaga | *Moringa olifera* | Jalgaon city, India | 14 | 25.5 | Unknown | 626.0 | 789.0 | **370.8** | Suryawanshi et al. 2014 |
| Neem | *Azadirachta indica* | Jalgaon city, India | 50 | 22.3 | Unknown | 487.0 | 614.0 | **288.6** | Suryawanshi et al. 2014 |
| Gulmohar | *Delonix regia* | Jalgaon city, India | 13 | 22.3 | Unknown | 486.0 | 612.0 | **287.6** | Suryawanshi et al. 2014 |
| Subabhule | *Acacia nilotica* | Jalgaon city, India | 32 | 19.1 | Unknown | 367.0 | 462.0 | **217.1** | Suryawanshi et al. 2014 |
| Sissam | *Dalbergia sisso* | Jalgaon city, India | 22 | 19.1 | Unknown | 286.0 | 360.0 | **169.2** | Suryawanshi et al. 2014 |
| Pilmohar | *Peltaforum pterocarpum* | Jalgaon city, India | 17 | 19.1 | Unknown | 380.0 | 479.0 | **225.1** | Suryawanshi et al. 2014 |
| Palas | *Butea monosperma* | Jalgaon city, India | 15 | 12.7 | Unknown | 141.0 | 178.0 | **83.7** | Suryawanshi et al. 2014 |
| Shirish | *Albizia lebbek* | Jalgaon city, India | 27 | 10.6 | Unknown | 96.0 | 121.0 | **56.9** | Suryawanshi et al. 2014 |
| Nilgiri | *Eucalyptus citriodora* | Jalgaon city, India | 28 | 9.6 | Unknown | 72.0 | 91.0 | **42.8** | Suryawanshi et al. 2014 |
| Teak | *Tectona grandis* | Jalgaon city, India | 244 | 9.6 | Unknown | 76.0 | 96.0 | **45.1** | Suryawanshi et al. 2014 |
| Amarillon, Roble | *Terminalia amazonia* | La Selva, Costa Rica | 547 | 22.3 | 13 years | N/A | 314.3 | **147.7** | Redondo-Brenes et al. 2006 |
| Cedro Maria | *Callophyllum brasiliense* | La Selva, Costa Rica | 612 | 20.1 | 13 years | N/A | 213.7 | **100.4** | Redondo-Brenes et al. 2006 |
| Almendro | *Dipteryx panamensis* | La Selva, Costa Rica | 752 | 14.7 | 13 years | N/A | 174.2 | **81.9** | Redondo-Brenes et al. 2006 |
| Chancho, Mayo | *Vochysia guatemalensis* | La Selva, Costa Rica | 664 | 25.5 | 13 years | N/A | 161.8 | **76.0** | Redondo-Brenes et al. 2006 |
| Fruta Dorada | *Virola koschnyi* | La Selva, Costa Rica | 840 | 23.8 | 13 years | N/A | 121.6 | **57.2** | Redondo-Brenes et al. 2006 |
| Pilon | *Hyeronima alchorneoides* | La Selva, Costa Rica | 771 | 16.5 | 12 years | N/A | 116.7 | **54.8** | Redondo-Brenes et al. 2006 |
| Botarrama | *Vochysia ferruginea* | La Selva, Costa Rica | 723 | 22.5 | 12 years | N/A | 104.8 | **49.3** | Redondo-Brenes et al. 2006 |
| Jacaranda | *Jacaranda copaia* | La Selva, Costa Rica | 684 | 22.6 | 13 years | N/A | 75 | **35.3** | Redondo-Brenes et al. 2006 |
| Ajillo | *Balizia elegans* | La Selva, Costa Rica | 781 | 19.0 | 12 years | N/A | 67.3 | **31.6** | Redondo-Brenes et al. 2006 |

\*Measured as basal diameter 2.5 cm above-ground (this study) or as diameter at breast height (dbh). #Above-ground biomass was estimated using an allometric equation for dry tropical conditions (Chave et al. 2005; Spring Gully and St Lucia) or was directly measured (Gatton). <sup>‡</sup>dbh measurements were converted into biomass (kg) using the allometric equation (Chave et al. 2005). <sup>^</sup>593 out of 680 trees survived to 3.5 years at the Spring Gully trial site.

Mature Pongamia trees growing in various locations around Brisbane exhibit remarkable growth despite little-to-no management, such as an 11 year-old tree in the suburb of St Lucia with an estimated total biomass of 902.6 kg and an organic carbon (CO_2_) content of 540.9 (1983.3) kg (Table 2). The considerable growth exhibited by individual trees planted for ornamental appeal, rather than in plantations, can be partially explained by the absence of competition for sunlight, water and resources from neighbouring trees.

Significant variation in the carbon sequestration potential of other tropical – subtropical trees has been reported for a range of species (Table 2). Age, environment, density and genetics were also attributed to differences observed. In Jalgaon, India, *Moringa olifera*, *Azadirachta indica*, and *Delonix regia* were among the top-performing species, sequestering 370.8, 288.6, and 287.6 kg of carbon per tree, respectively (Suryawanshi et al., 2014) (Table 2). An assortment of twelve to thirteen year-old trees growing at La Selva, Costa Rica, also exhibited varying carbon sequestration potentials, ranging from 31.6 to 147.7 kg per tree (Redondo-Brenes and Montagnini, 2006) (Table 2).

### Assessing Drought Tolerance in Pongamia

Pongamia is reported to be highly drought-tolerant (Degani et al., 2022), making it suitable for growth in low rainfall areas, including for capturing carbon in arid, drought-prone regions. To further assess drought tolerance, Pongamia seedlings were grown in the glasshouse and supplied with either a well-watered or restricted watering regime (Figure 8). Weekly growth measurements were taken on total fresh/dry weight, total turgid weight and shoot/root length. Relative water content was calculated using the difference between fresh and dry weight over the difference between turgid and dry weight.

**Figure 8.**
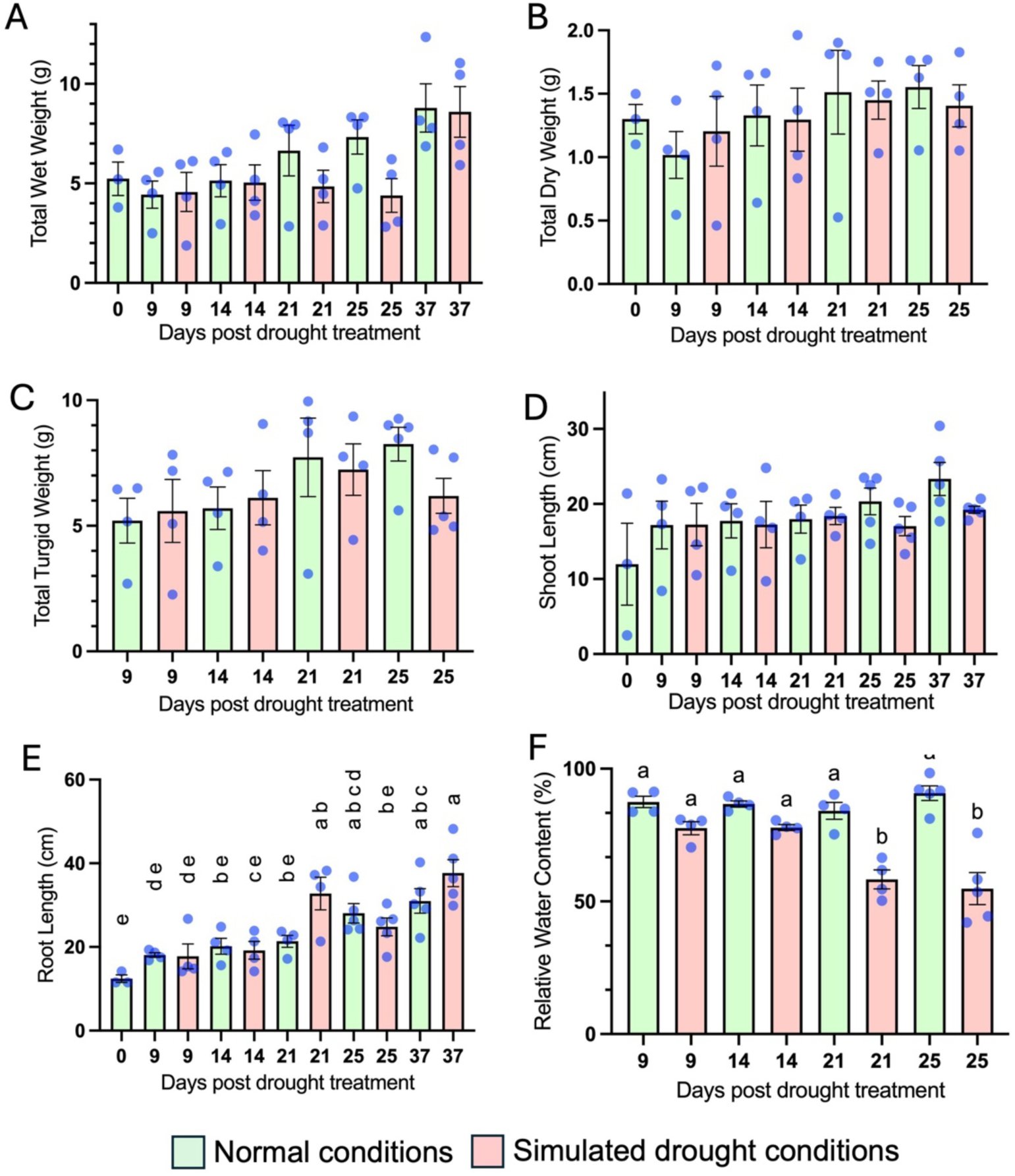
Pongamia drought experiment. Pongamia plants were subjected to well-watered or restricted watering regimes. **(A)** Total fresh weight, **(B)** total dry weight, **(C)** total turgid weight, **(D)** shoot length, **(E)** root length and **(F)** relative water content expressed as a percentage using the formula RWC=[(FW−DW)/(TW−DW)]×100, were recorded at various time points. Numbers indicate days post drought treatment. Letters indicate separation of means into homogeneous sets by post-hoc Tukey test at p ≤ 0.05. For each time point, n = 3 to 5 biological replicates.

Relative water content (RWC) reflects how much water a plant is holding relative to its fully hydrated state. It is often used to assess a plant’s water status and ability to cope with limited water availability, providing insights into its drought tolerance. Plants with higher RWC under drought conditions are better adapted to drought, as they can retain water more effectively in their tissues. In this study, the RWC of drought-treated plants remained high for the first 3 weeks of the experiment and were significantly reduced to 50% at 21-25 days of the final week (Figure 8F). This suggests that Pongamia can generally withstand drought conditions and maintain water content and cellular function under drought stress.

No significant differences in shoot length were observed during the experiment (Figure 8D), although this would be anticipated over time. Significant differences in root lengths were observed, with average root length of plants grown under drought being longer than the controls at 21 and 37 days (Figure 8C). This could indicate that the roots of the plants subjected to drought were elongating in an attempt to search for water.

## DISCUSSION

Pongamia is a fast-growing tropical legume tree that can accumulate biomass quickly. This is ideal for the sequestration of carbon. Here, we demonstrate that trees in the sub-optimal growing region of Spring Gully, Australia, are estimated to have sequestered 2.89 t C/ha (10.59 t CO_2_/ha) within 3.5 years when grown at a density of 680 trees per 1.5 ha. In the more optimal growing region of Gatton, Australia, Pongamia trees were estimated to sequester 4.08 t C/ha (14.96 t CO_2_/ha) within 4.5 years at a density of 450 trees per ha. These figures are comparable to a previous report from Singrauli, India (Bohre et al., 2014), where 4 year-old trees were estimated to sequester 2.18 t C/ha (7.99 t CO_2_/ha). Indeed, the potential for Pongamia to sequester carbon is reported to be greater than for many other tree species (Prasad, 2022). Mature Pongamia trees were found to sequester approximately 45-50 kg of carbon per annum, which compares extremely favourably to other trees such as Neem (28-35 kg), Mahua (23-26 kg), and Tendu (11-15 kg) (Degani et al., 2022). Moreover, Pongamia trees aged over 20 years, can sequester 15-25 tonnes carbon per annum per hectare, with individual Pongamia trees having the potential to sequester over 700 kg of carbon over a period of 25 years (Degani et al., 2022). Collectively, the data highlight Pongamia’s strong potential for sequestering carbon. Additional biomass data collected from older Pongamia trees of different genetics, growing in varying climatic regions under different management practices would further strengthen our understanding of this potential and enable more accurate predictions of their ability to sequester carbon throughout their lifespan.

Differences in carbon sequestration of Pongamia between locations can be explained by classical Genotype by Environment by Management (G x E x M) interactions (Annicchiarico, 2002). In Spring Gully and Gatton, biomass differences observed within trees from same mother are likely due to genetics (Gresshoff et al., 2015), as the environment was roughly uniform within each site. This is because Pongamia is an outcrossing species and progeny seed from a single mother tree can be highly heterozygous (and thus phenotypically heterogeneous), as confirmed by DNA fingerprinting using molecular markers (Jiang and Gresshoff, 1997). Therefore, plants raised from seed, such as those planted at Spring Gully and Gatton, are genetically diverse and can exhibit variations in traits such as height, biomass, canopy structure, root architecture, seed morphology, oil content and yield (Sharma et al., 2011; Murphy et al., 2012). Such variation can be reduced by growing clonal material derived from cuttings or through tissue culture, but those strategies can be slower and more labour intensive. Whether trees derived from these strategies differ in their carbon sequestration potential as they mature is unknown.

The carbon sequestering potential of trees is highly dependent on where and how the trees are planted. Often the lack of preliminary planning can lead to low survival and growth rates of trees (Cao, 2011; Cao et al., 2011; Xu, 2011). Similarly, planting time can have a considerable impact on plant establishment. Saplings planted at Spring Gully in May (*i.e.,* late autumn in Australia) were exposed to frost events shortly after, which stifled growth and led to some dieback as the trees had insufficient time to harden-off and develop adequate biomass to withstand the frost. Planting areas should also be carefully selected to have sufficient slope and air circulation to restrict frost (Wylie et al., 2023). A machine learning framework that integrates environmental variables to quantify tree growth and its relationship with tree numbers have been used to assess biomass and carbon sequestration potentials (Yao et al., 2024). The tree planting potential of an area can be determined by its historical tree cover, where an area once extensively covered by forests is more likely to provide strong tree growth.

Carbon sequestration and restoring diversity to grassland ecosystems or the restoration of marginal lands can contribute to climate change mitigation via carbon sequestration (De Deyn et al., 2011; Gelfand et al., 2013; Nackley et al., 2013; Awasthi et al., 2017). Agriculturally degraded or abandoned lands also have the potential to sequester atmospheric CO₂ as soil organic matter during natural succession (Yang et al., 2019), though this process can be slow, requiring extended periods to restore soil carbon levels to their pre-agricultural state. C4 grasses and legumes, such as Pongamia, are particularly well-suited for this purpose due to their adaptability to degraded lands and their efficient photosynthetic mechanisms or nitrogen-fixing capabilities, respectively (Yang et al., 2019). Pongamia also represents an ideal candidate for phytoremediation of lands that are unusable for agriculture, such as mine restoration sites. This is due to its tolerance to drought, salinity, and heavy metals in the soil (Degani et al., 2022), as well as its ability to form symbiotic associations with beneficial soil microbes, such as rhizobia and mycorrhiza, that provide the tree with critical nutrients. Moreover, Pongamia has potential to exclude and detoxify heavy metals in root issues by translocating and retaining contaminants in above-ground biomass (Kumar et al., 2017). Preliminary results from an on-going Pongamia plantation on a copper mine site in Zambia reveal high survival rates of Pongamia trees under elevated soil copper concentrations of up to 6,000 mg/kg (Warr, 2019).

The early establishment of plantations, ensuring survival of the seedlings with maximum growth and carbon sequestration potential, can be limited by water availability. For this reason, drought tolerance can be a highly beneficial trait. Our findings demonstrate that even very young Pongamia seedlings exhibit drought tolerance. Irrigation has been deemed necessary during seedling establishment of some Pongamia plantations (Murphy et al., 2012; Degani et al., 2022). However, much more work is needed to understand optimal G x E x M combinations for Pongamia, with very few studies having explored adaptive mechanisms of drought tolerance in the legume tree (Rex Immanuel et al., 2019). As Pongamia trees mature, their dense network of lateral and tap roots, reported to extend up to 9-10 m below ground (Rex Immanuel et al., 2019), also helps to confer drought tolerance. This further supports their ability to rapidly accumulate biomass and sequester carbon, even in low rainfall years. This also enables Pongamia trees to survive long periods of drought (Murphy et al., 2012). Further studies comparing irrigation at establishment and at maturity are needed to better understand how water availability impacts biomass accumulation, carbon sequestration and seed yield.

## CONCLUSION

In this study, the growth characteristics and carbon sequestration potential of *Pongamia pinnata* trees grown in subtropical environments were assessed. Pongamia was found to have considerable potential to sequester carbon, even when grown in substandard growing conditions. Its tolerance to heavy metals, drought, and frost, as well as its ability to form symbiotic associations with beneficial mycorrhizal fungi and nitrogen-fixing rhizobia bacteria, further underscores the potential for growing Pongamia on marginal lands, including those used for land restoration, and providing social and economic benefits. This study also underscores the importance of pursuing additional, longer term research initiatives to establish and optimise Pongamia’s potential to sequester carbon over extended periods. This includes evaluating plantations of different feedstock genetics, at different locations (varying in soil, climate, etc.), and different management regimes (planting density, pruning, irrigation, fertiliser, symbiont inoculation, etc.).

## MATERIALS AND METHODS

### Spring Gully Plantation

At Spring Gully, near Roma, Queensland, 680 Pongamia trees were planted in 2008 across roughly 1.5 ha. Seedlings were germinated from seed collected from six genetically distinct mother stocks of Pongamia located around Brisbane. The seedlings were established in pots in an evaporatively-cooled glasshouse (∼28°C) and planted at the Spring Gully trial site when they were roughly 50 cm tall. Individual trees were planted 4 m apart in 20 rows that were spaced 5 m apart. Following planting, the seedlings were irrigated manually using water-tanks fitted on the back of a truck. The water was supplied through a desalination plant located on site. Pipe irrigation was also established and used on a need basis.

### Gatton Plantation

At the University of Queensland, Gatton campus field site, 450 Pongamia trees were planted in December 2008 across roughly 1 ha. The seedlings were derived from seed collected from trees located around Brisbane. They were initially germinated in pots in an evaporatively-cooled glasshouse (∼28°C) located at the University of Queensland, St Lucia campus. The seedlings were an average of ∼50 cm tall when planted, and were spaced 0.6-0.7 m apart across three rows, each approximately 100 m long, with 9 m between each row. The plot was irrigated using in-ground drip lines and watered on a need basis. At subsequent intervals after planting, destructive sampling was performed on four to six trees to obtain biomass measurements (*e.g.,* the weight of the trunk, woody tissues, root system, leaf, and pruned foliage). Plant materials were dried in an oven at ∼60°C for approximately 5 days to determine the constant dry weight.

### Soil Component Analysis

Soil analyses were performed at the Analytical Services Unit of the University of Queensland, St Lucia campus. Extraction procedures are outlined in (Rayment and Higginson, 1992).

### Drought Analysis

Seedlings established from seed were grown in ANOVA pots containing UQ23 mix planted in an evaporatively-cooled glasshouse (∼28°C) located at the University of Queensland, St Lucia campus. Each treatment group (control and drought) consisted of 22-25 plants. The plants in the control group were watered at a 3-day interval whereas the plants subject to drought did not receive additional water for the duration of the experiment. The soil was sufficiently watered for both groups at the start of the experiment. At each time point, plants were destructively sampled for measuring the wet and turgid weight, as well as shoot and root length. Plant material was subsequently dried in an oven at 60°C to obtain the dry weight.

## ACKNOWLEDGMENTS

Present and former colleagues are thanked for their contributions and discussions over the years. Origin Energy and Hewitt are thanked for providing the land at Spring Gully. George Muirhead (BPA Pty Ltd) is thanked for extensive collaboration for the Spring Gully field trial. This work was supported by funding from the Australian Research Council Centre of Excellence scheme (CEO348212) and from Rio Tinto. We also thank Dr Joe Eyre for his feedback and suggestions on the manuscript.

## Notes

### Competing Interest Statement

The authors have declared no competing interest.

