## Supplemental Figure 1, Supplemental Table 1, Supplemental Table 2. for "The biofuel producing legume tree, *Pongamia pinnata*, offers strong potential for carbon sequestration"

[2] Current address: Terragen Biotech Pty Ltd, Coolum Beach QLD 4573

ORCID IDs:

ANDY CHEN: 0000-0002-4004-1283

PAUL SCOTT: 0009-0000-4993-9690

PETER GRESSHOFF: 0000-0003-0693-2813

BRETT FERGUSON: 0000-0003-3737-8368

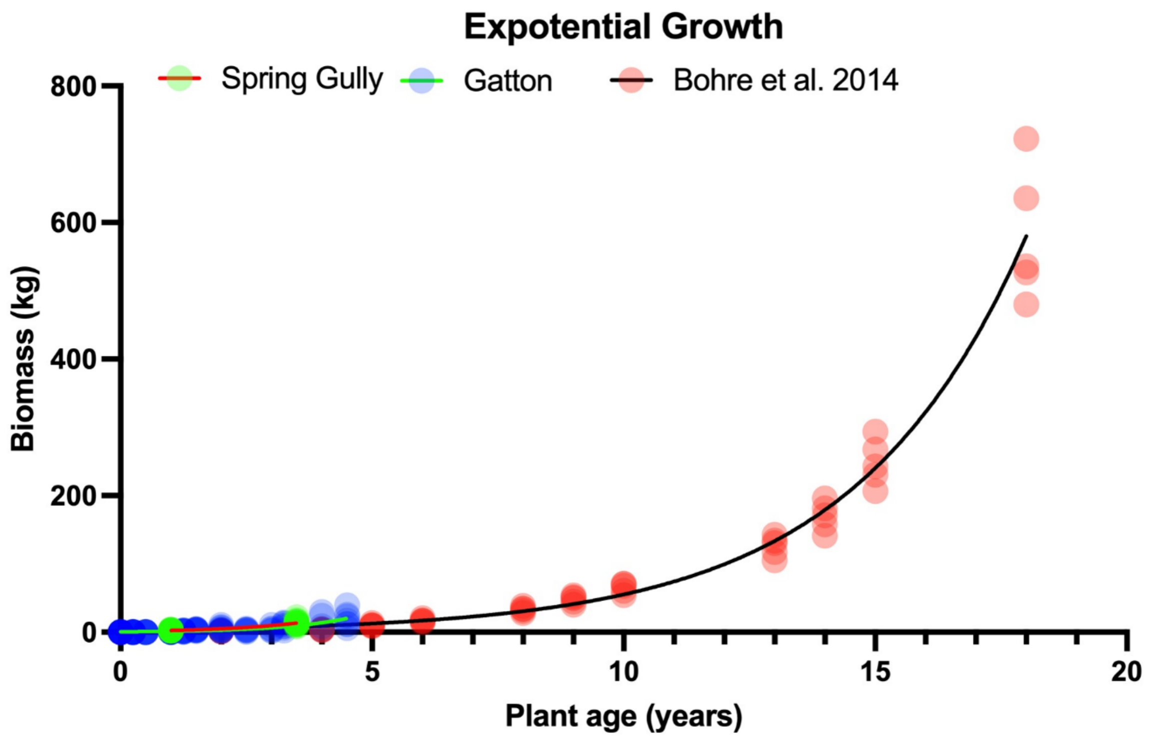


Supplemental Figure 1.

Biomass per tree predicted using an exponential (Malthusian) growth model. Exponential growth was modelled as Y=Y0*exp(k*x), where Y is biomass, Y0 is the starting biomass, K is the Rate constant for growth, and x is time. Goodness of Fit showed an R squared value of 0.67 for the data from Gatton and 0.97 for the data derived from Bohre et al. (2014).

Supplemental Table 1. Genotype of Pongamia seedlings planted in each row of the Spring Gully plantation.

| Row | Genotypes planted | Number of trees planted |
|---|---|---|
| 1 | "Tree 8" | 8 |
| 1 | "Tree 29" | 27 |
| 2 | "Tree 29" | 34 |
| 3 | "Tree 29" | 34 |
| 4 | "Tree 29" | 34 |
| 5 | "Tree 29" | 34 |
| 6 | "Tree 29" | 34 |
| 7 | "Tree 29" | 34 |
| 8 | "Tree 29" | 17 |
| 8 | "Tree 19" | 17 |
| 9 | "Tree 19" | 34 |
| 10 | "Tree 19" | 34 |
| 11 | "Tree 19" | 34 |
| 12 | "Tree 19" | 22 |
| 12 | "Tree 17" | 12 |
| 13 | "Tree 17" | 34 |
| 14 | "Tree 17" | 13 |
| 14 | "Tree 20" | 21 |
| 15 | "Tree 20" | 34 |
| 16 | "Tree 20" | 34 |
| 17 | "Tree 20" | 16 |
| 17 | "Tree 22" | 18 |
| 18 | "Tree 22" | 34 |
| 19 | "Tree 22" | 34 |
| 20 | "Tree 22" | 34 |

Supplemental Table 2. Nutrients and trace elements detected in water samples collected from the Spring Gully Pongamia trial site.

| Trace elements | Concentration | Sample 1 | Sample 2 | Sample 3 |
|---|---|---|---|---|
| Al | mg/L | 0.028 | 0.005 | 0.001 |
| As | mg/L | 0.023 | 0.008 | 0.002 |
| Ca | mg/L | 1.121 | 0.998 | 0.960 |
| Cd | mg/L | 0.002 | 0.002 | 0.000 |
| Co | mg/L | 0.002 | 0.010 | 0.001 |
| Cr | mg/L | 0.010 | 0.008 | 0.007 |
| Cu | mg/L | 0.004 | 0.000 | 0.000 |
| Fe | mg/L | 0.008 | 0.003 | 0.000 |
| Hg | mg/L | 0.002 | 0.001 | 0.001 |
| K | mg/L | 0.756 | 0.714 | 0.778 |
| Mg | mg/L | 0.049 | 0.017 | 0.002 |
| Mn | mg/L | 0.001 | 0.002 | 0.001 |
| Na | mg/L | 19.661 | 19.376 | 19.138 |
| Ni | mg/L | 0.005 | 0.002 | 0.000 |
| P | mg/L | 1.147 | 0.761 | 0.563 |
| Pb | mg/L | 0.001 | 0.002 | 0.004 |
| S | mg/L | 0.031 | 0.027 | 0.002 |
| Se | mg/L | 0.004 | 0.002 | 0.002 |
| Zn | mg/L | 0.003 | 0.000 | 0.002 |
